# Experimental and simulated CO_2_ responses of photosynthesis in leaves of *Hippophae rhamnoides* L. under different soil water conditions

**DOI:** 10.1101/838284

**Authors:** Qin Wu, Cheng Li, Qiang Chen

## Abstract

CO_2_ concentrations and soil moisture conditions seriously affect tree growth and physiological mechanisms. CO_2_ responses of photosynthesis are an important part of plant physiology and ecology research. This study investigated the photosynthetic CO_2_ responses in the leaves of two-year-old *Hippophae rhamnoides* L. under eight soil water conditions in a semi-arid loess hilly region, and discussed the quantitative relationship between CO_2_ responses and soil moisture. CO_2_ response curves and parameters were fitted using a rectangular hyperbola model, non-rectangular hyperbola model, exponential equation, and modified rectangular hyperbola model. Results revealed that the relative soil water content (*RWC*) required to maintain a high photosynthetic rate (*P*_n_) and carboxylation efficiency (*CE*) ranged from 42.8% to 83.2%. When *RWC* fell outside these ranges, the photosynthetic capacity (*P*_nmax_), *CE*, and CO_2_ saturation point (*CSP*) decreased. CO_2_ response curves and three parameters, *CE*, CO_2_ compensation point (*Γ*), and photorespiration rate (*R*_p_), were well fitted by the four models when *RWC* was appropriate. When *RWC* exceeded the optimal range, only the modified rectangular hyperbola model fitted the CO_2_ response curves and photosynthetic parameters better.

## Introduction

Photosynthesis is a complex process affected by many factors in plants, including CO_2_ and water, which have important effects[1,2]. CO_2_ is the substrate of photosynthesis, and its atmospheric concentration is predicted to reach ~550 μmol·mol^−1^ by 2050[3,4]. Global water shortages are aggravated by changes due to increasing CO_2_ concentrations and the warming climate[5,6]. The increase in CO_2_ can cause global climate change and directly affect the metabolism and growth of plants[7,8]. Drought affects plant growth and development severely [9,10], as well as limits photosynthesis through carbon metabolism by restricting CO_2_ diffusion[11,12]. However, plants display adaptability and resistance to water deficits[13,14]. Moreover, photosynthetic efficiency is higher within a certain water range, which varies according to the plant species and photosynthetic mechanism[15,16]. CO_2_ responses are an important part of plant physiology and ecology research, the measurement and simulation of which are the main approaches for studying photosynthesis[17,18]. The photosynthesis CO_2_ response model has played an important role in increasing our understanding of photosynthetic carbon uptake, which has thereby improved our understanding and predictions of plant photosynthetic physiology and its response to environmental changes and biogeochemical systems[19,20]. CO_2_ response curves reflect the quantitative relationship between plant net photosynthetic rate (*P*_n_) and CO_2_ concentration and can be used to estimate photosynthetic parameters, including the CO_2_ saturation point (*CSP*), photosynthetic capacity (*P*_nmax_), compensation point (*Γ*), carboxylation efficiency (*CE*), and photorespiration rate (*R*_p_)[21,22].

CO_2_ responses have been fitted using biochemical models[23], empirical models[24,25], and optimized models[26,27], which are based on biochemical models. Biochemical models calculate two key model parameters, the maximum rate of carboxylation (*V*_cmax_) and the maximum electron transport rate (*J*_max_)[28,29]. Empirical models include the Michaelis-Menten model [30], rectangular hyperbola model[31], non-rectangular hyperbola model [32], and exponential equation [33], which have been applied in most crops[34,35]and some woody species[36,37]. Ye [38]thought the Michaelis-Menten and rectangular hyperbolic models were essentially the same. In recent years, some studies have proposed an improved rectangular hyperbolic model, namely, the modified rectangular hyperbola model[39,40]. This model has been applied to some plants, including some gramineous plants[41,42], herbs[43,44], and woody plants[45,46]. Results revealed that this new model could overcome the limitations of traditional models and accurately fit the CO_2_ response curve and its characteristic parameters. Previous studies on photosynthesis CO_2_ response models have focused on the estimation and optimization of key parameters in field crops[47,48]. However, the applicability of different models simulating the CO_2_ response data of woody plants under adverse conditions, such as continuous drought, has rarely been reported.

*Hippophae rhamnoides* L. is a common afforestation species found in the arid and semi-arid regions of Northern China, which has a high economic value and plays an important role in ecological restoration and soil and water conservation. *H. rhamnoides*L. is a non-leguminous and nitrogen-fixing species, deciduous shrubs, and is resistant to barren and dry conditions. In recent years, studies have focused on its growth[49], water consumption[50,51], and photosynthetic light response characteristics[52,53]. These studies have been conducted under water stress, while only a few studies related to the physiological characteristics of drought stress have been conducted[54,55]. However, continuous observations and the examination of photosynthesis CO_2_ response in leaves of *H*. *rhamnoides* L. at many soil moisture gradients during the accelerated soil drought process have not been addressed. Therefore, the quantitative relationship between the photosynthetic CO_2_ response process and soil moisture remains unclear.

In this study using potted seedlings of *H*. *rhamnoides* L., CO_2_ response curves and parameters were evaluated and fitted with the rectangular hyperbola model, non-rectangular hyperbola model, exponential equation, and modified rectangular hyperbola model under different soil moisture conditions. The goals of this study were to define the quantitative relationship between photosynthetic CO_2_ response processes and soil moisture, as well as explore the applicability of different CO_2_ response models to fit CO_2_ response processes and parameters. The findings of this study will provide an in-depth understanding of the photosynthetic physiology-ecological characteristics and cultivation of *H*. *rhamnoides* L. in the loess hilly-gully region of Northern China. Furthermore, the applicability of different CO_2_ response models can be evaluated from these findings and used in future studies.

## Material and methods

### Study area

The experimental site was located in the Tuqiaogou watersheds (37°36′58″N, 110°02′55″E) of Yukou Town, Fangshan County, Shanxi Province, China, a portion of the gully-hilly area of the Loess Plateau in the middle reaches of the Yellow River. This area has a sub-arid, warm temperate, continental monsoon climate. The average annual precipitation is 525.0 mm with more than 70% of the precipitation concentrated between July and September. The annual potential evaporation is 1839.7 mm with the greatest amount of evaporation occurring between April and June. The annual frost-free period lasts 140 d. The soil is classified as medium loessial soil, and the soil texture is uniform with a pH value ranging from 8.0 to 8.4. Vegetation consists mainly of trees, shrubs, lianas, and subshrubs. Tree species are predominantly *Robinia pseudoacacia*, *Ulmus pumila*, *Platycladus orientalis*, and *Syringa oblata*. Shrubs are mainly *Rosa xanthina* and *Ulmus macrocarpa*. Herbs consist of *Compositae* and *Gramineae*, of which the *Compositae* belong to the *Artemisia* genus. Most of the forest land consists of sparse woodland with poor stand stability.

### Materials and water treatments

Two-year-old *H*. *rhamnoides* L. were used as the experimental materials and were selected carefully to ensure consistency in their height, diameter, and growth. Plants were investigated and marked one by one before transplantation. In March 2018, seedlings were transplanted in containers (50 cm in height, 35 cm in diameter) that had drainage holes in the bottom. A total of six basins with one plant in each pot were used. The relative soil water content (*RWC*) and photosynthetic CO_2_ responses in the leaves of *H*. *rhamnoides* L. were determined in August. Three strong plants were selected and watered to saturation, and the initial *RWC* was obtained; the first CO_2_ response was also determined. Then, soil moisture gradients were obtained every two days through the natural water consumption method after artificially supplying water. The soil mass water content (*MWC*, %) was measured by the stoving method. The *RWC* was considered as the ratio of *MWC* to the field water capacity (*FC*, %). The potting soil *FC* was 24.3%, according to the cutting ring method, and the soil bulk density was 1.26 ± 0.13 g**·**cm^−3^. Eight *RWC* gradients were obtained and found to be 91.7%, 83.2%, 71.5%, 54.6%, 42.8%, 31.9%, 26.1%, and 21.4%. The experiment was monitored under a canopy with a plastic film covering the top on rainy days to prevent rain from interfering with the *RWC*.

### CO_2_ response determination

Three strong, mature leaves were selected and marked in a central test plant. CO_2_ responses under different soil moisture conditions were measured using a CIRAS-2 (PP Systems, Hitehin, UK) portable photosynthesis system. The light saturation point for *H. rhamnoides* L. was 1400 μmol**·**m^− 2^ **·**s^−1^ [52,54]. Measurements were obtained under each soil moisture condition on separate days. The time of measurements occurred from 08:30 to 11:00 h in completely clear weather to reduce the effects of outside light fluctuations. Measurements were obtained three times for each leaf, and the average value was calculated and used for the analyses. The atmospheric temperature ranged from 24°C to 26°C, and the relative humidity was approximately 60% ± 4.0%. The CO_2_ concentration in the leaf chamber was controlled and regulated from 0 to 1400 μmol**·**mol^−1^ by a small cylinder with high CO_2_ concentrations. The CO_2_ concentration gradients were 1400, 1200, 1000, 800, 600, 400, 200, 180, 150, 120, 90, 60, 30, and 0 μmol**·**mol^−1^. The duration of the measurement lasted 120 s at each CO_2_ concentration, and the apparatus automatically recorded the photosynthetic physiological parameters, including the *P*_n_ (μmol·m^−2^·s^−1^) and intercellular CO_2_ concentration (*C*_i_, μmol·mol^−1^).

### Data analysis

CO_2_ response curves were drawn with *C*_i_ as the horizontal axis and *P*_n_ as the vertical axis. According to the measured data point trends, *CSP* (μmol·mol^−1^), *P*_nmax_ (μmol·m^−2^·s^−1^), and *Γ* (μmol·mol^−1^) were estimated and regarded as measured values. *CE*_Γ_ (mol·m^−2^·s^−1^) at *Γ*, the intrinsic carboxylation efficiency (*CE*_0_, mol·m^−2^·s^−1^) at 0 *Γ*, the absolute value (*CE*_Γ0_, mol·m^−2^·s^−1^) of the slope of the line from *C*_*i*_ = 0 to *C*_*i*_ = *Γ* in the CO_2_ response curve, and *R*_p_ (μmol·m^−2^·s^−1^) were calculated using the traditional linear regression method and used as the measured values to compare to the fitted values of the four models.

Statistical analyses were performed using Microsoft Excel 2003(Microsoft Corp., Redmond,Wash.). Significant differences were analyzed by a one-way ANOVA and Duncan’s post-hoc test. Nonlinear regression was analyzed using SPSS v18.0 (SPSS Inc., Chicago, Illinois). Data were expressed as the mean ± standard deviation (S.D.), and significance was interpreted as *p* < 0.05. The CO_2_ response curve was fitted using the rectangular hyperbola model, non-rectangular hyperbola model, exponential equation, and modified rectangular hyperbola model (described below).

### Rectangular hyperbolic model

The rectangular hyperbolic model is expressed as follows[31]:

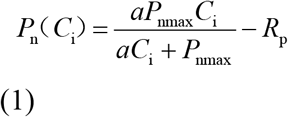

where *P*_n_ is the net photosynthesis rate, *C*_i_ is the intercellular CO_2_ concentration, α is the slope of the CO_2_ response curve when *C*_i_ = 0 (namely, the initial slope of the CO_2_ response curve and the initial *CE*), *P*_nmax_ is the photosynthetic capacity, and *R*_p_ is the photorespiration rate.

*CE*_Γ_, *CE*_0_, and *CE*_Γ0_ are expressed as follows:

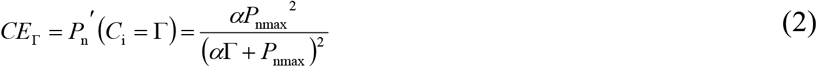

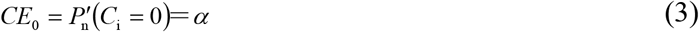

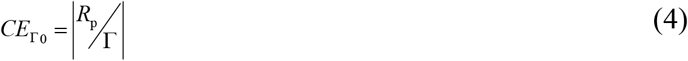

*Γ* is expressed as follows:

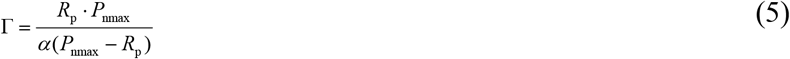

where the line y = *P*_nmax_ intersects the linear equation when *C*_i_ is below 200 μmol mol^−1^, and the value of the intersected point on the x-axis is *CSP*[56].

### Non-rectangular hyperbola model

The non-rectangular hyperbola model is expressed as follows [32]:

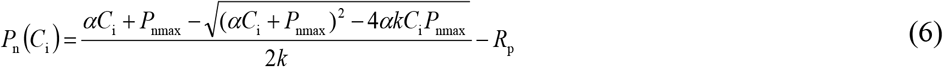

where *k* is the curved angle of the non-rectangular hyperbola; the definitions of other parameters are the same as above.

*CE*_Γ_, *CE*_0_, and *CE*_Γ0_ are expressed as follows:

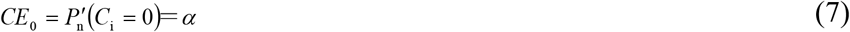

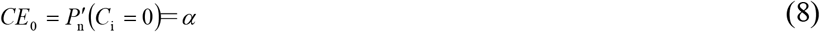

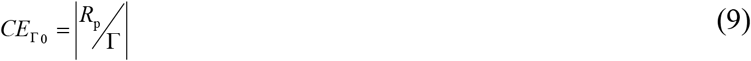

*Γ* is expressed as follows:

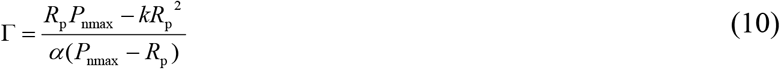

where the line y = *P*_nmax_ intersects the linear equation when *C*_i_ is below 200 μmol·mol^−1^, and the value of the intersected point on the x-axis is *CSP*[38].

### Exponential equation

The exponential equation is expressed as follows [33]:

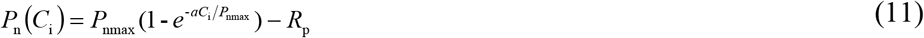

where the definitions of *P*_n_, *C*_i_, *P*_nmax_, α, and *R*_p_ are the same as above.

*CE*_Γ_, *CE*_0_, and *CE*_Γ0_ are expressed as follows:

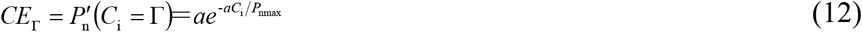

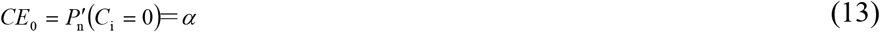

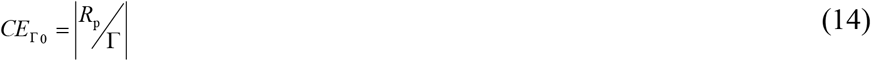

*Γ* is expressed as follows:

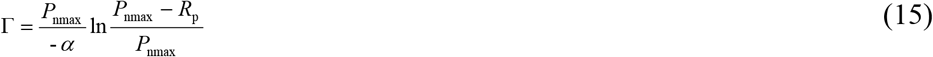

where the line y = *P*_nmax_ intersects with the linear equation at *C*i ≤ 200 μmol·mol^−1^, and the value of the intersected point on the x-axis is *CSP* [57].

### Modified rectangular hyperbola model

The modified rectangular hyperbola model is expressed as follows[39]:

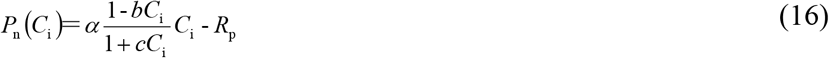

where b and c are coefficients; the definitions of other parameters are the same as above.

*CE*_Γ_, *CE*_0_, and *CE*_Γ0_ are expressed as follows:

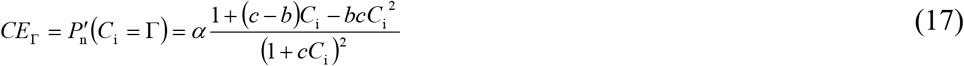

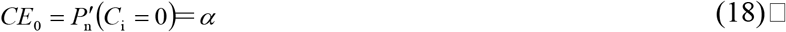

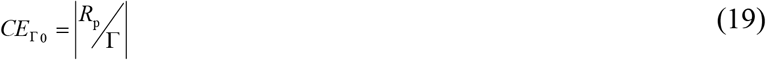

*CSP* and *P*_nmax_ are expressed as follows:

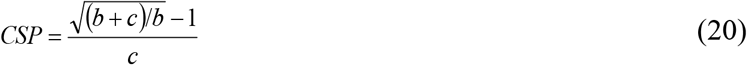

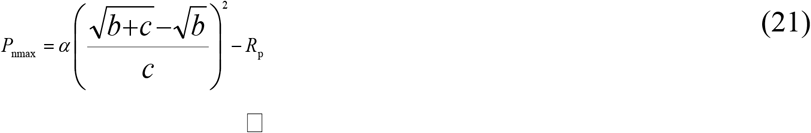

## Results

### Photosynthetic CO2 response

Soil moisture significantly affected the photosynthetic CO_2_ response of *H. rhamnoides* L. (Fig. 1). Under different soil moisture conditions, *P*_n_ increased rapidly as *C*_i_ increased, when *C*_i_ was below ~200 μmol·mol^−1^. *P*_n_ increased slowly as *C*_i_ increased, and the maximum *P*_nmax_ appeared at *CSP*. When *C*_i_ reached *CSP*, the CO_2_ response was considerably different under different soil water conditions, specifically when *RWC* ranged from 42.8% to 83.2%, *P*_n_ of each CO_2_ response curve changed slightly as *C*_i_ increased after *C*_i_ reached *CSP*. When *RWC* was out of the above ranges, *P*_n_ decreased considerably after *C*_i_ reached *CSP*, *P*_n_ in each curve at the highest *C*_i_ was significantly smaller than its *P*_nmax_ under the same soil moisture conditions (*p* < 0.05) (Table 1) Clearly, CO_2_-saturated inhibition had occurred. Furthermore, the CO_2_ responses to soil moisture had an obvious *RWC* threshold. The overall level of *P*_n_ in each CO_2_ response curve increased initially, then decreased as *RWC* decreased. The *P*_n_ level was the highest when *RWC* was 71.5%; thus, an increase or decrease in *RWC* led to a decrease in the overall *P*_n_ level. *CSP* and *P*_nmax_ were high and *P*_n_ did not decrease at high CO_2_ concentrations when *RWC* ranged from 42.8% to 83.2%; thus, these *RWC* ranges were suitable for photosynthesis in the leaves of *H. rhamnoides* L..

**Fig 1.** Photosynthetic CO_2_ response curves in the leaves of *H. rhamnoides* L. under different soil water conditions (mean ±S.D.).

**Table 1.**
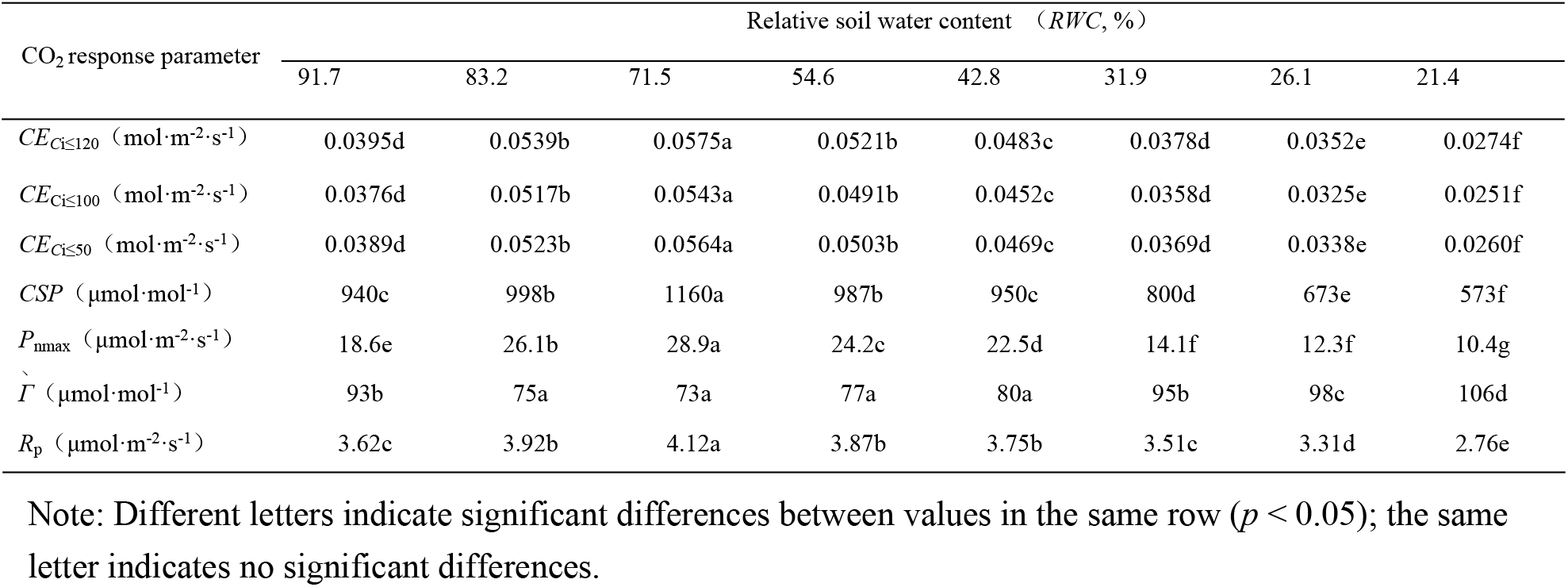
Data on the photosynthetic CO_2_ response parameters of *Hippophae rhamnoides* L. under different soil water conditions (mean ± S.D.).

### Simulation of CO_2_ response curves and characteristic parameters

The simulated effects of the four models fitting the CO_2_ response data were notably different under different soil moisture conditions (Fig. 2; Table 2). CO_2_ responses curves and photosynthetic characteristic parameters(*CE*_0、_*CE*_Γ_、*CE*_Γ_, *Γ*, and *R*_p_) were well simulated by the four models, and the determination coefficients were all > 0.991 when *RWC* ranged from 42.8% to 83.2% (Table 2). Moreover, within the above *RWC* range, *P*_nmax_ and *CSP* fitted by the modified rectangular hyperbola model were comparable with the measured value. The *P*_nmax_ values fit by the other three models were significantly higher than their observed values, while the fit *CSP* values were significantly lower than their observed values. When *RWC* was outside the range of 42.8%-83.2%, only the modified rectangular hyperbola model fit the CO_2_ responses (*R*^2^ > 0.99) and characteristic parameters well(Figue 2D; Table 2), the other three kinds of models produced large deviation to fit the CO_2_ response process and its characteristic parameters(Figues 2A,B,C; Table 2).

**Fig 2.**
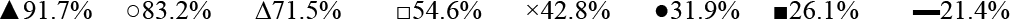
Photosynthetic CO_2_ response curves of the leaves of *Hippophae rhamnoides* L. simulated by four models under different soil water conditions (mean ± S.D.): A) Rectangular hyperbola model; B) Non-rectangular hyperbola model; C) Exponential equation; and D) Modified rectangular hyperbola model.

**Table 2.**
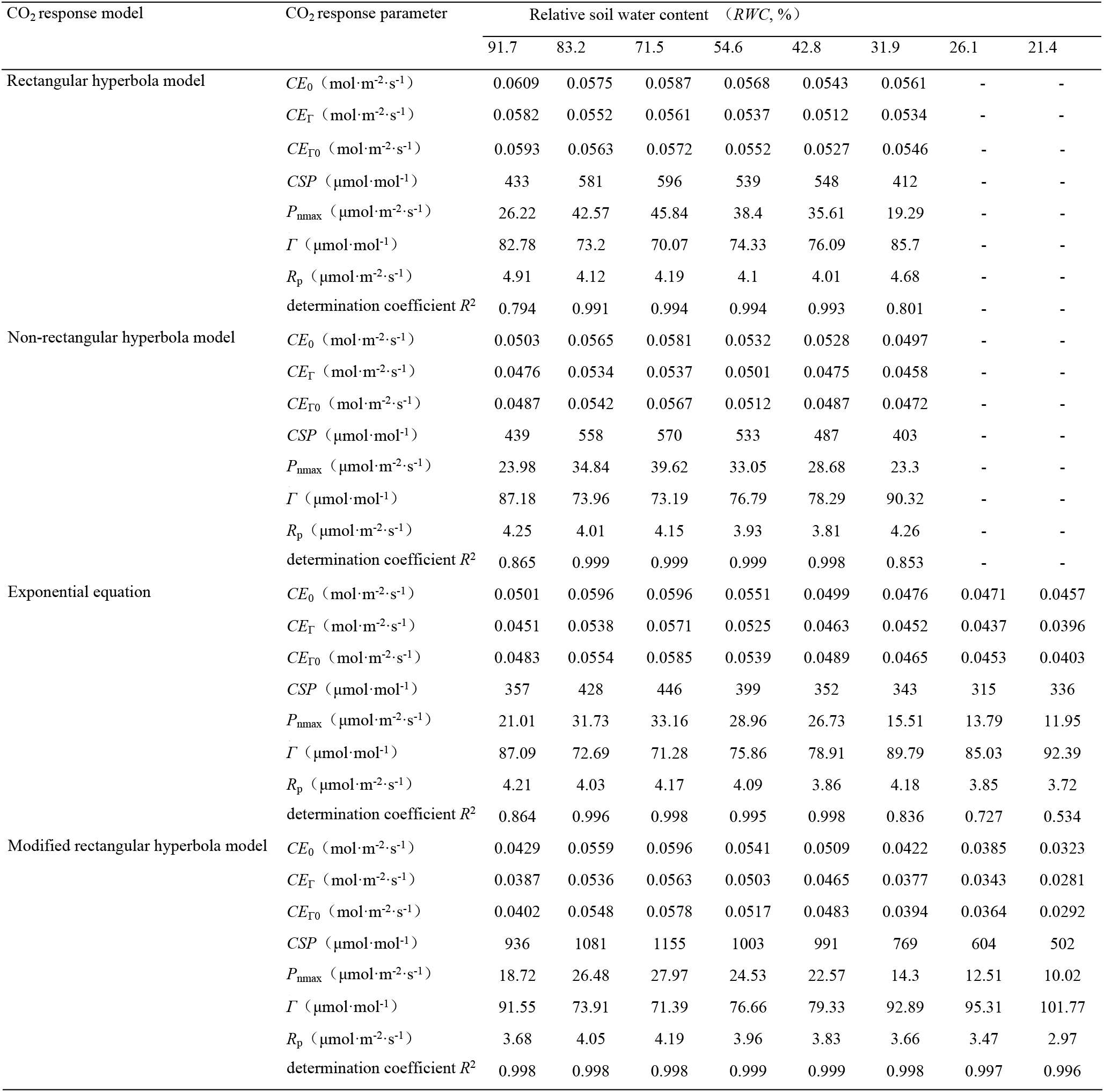
Data on the photosynthetic CO_2_ response parameters of *Hippophae rhamnoides* L. fitted by 4 models.

## Discussion

### Effects of soil moisture on CO_2_ response curves and photosynthetic parameters

Water is a major limiting factor in the recovery and restoration of vegetation found in the loess, hilly-gully regions of China. *RWC* seriously affected light-response curves and photosynthetic parameters, which also profoundly affected CO_2_ response curves and photosynthetic parameters in the leaves of *H. rhamnoides* L. The classical form of a *P*_n_-*C*_i_ curve can be summarized in three stages[58,59]. First, an approximately linear segment is observed when *C*_i_ ≤ 200 μmol·mol^−1^. Thus, *P*_n_ increases rapidly as *C*_i_ increases, namely, during the ribulose bisphosphate (RuBP) restriction phase. The slope of the straight line is the mesophyll conductance, *CE*, which reflects the assimilative capacity of plant responses to low CO_2_[60,61]. Second, the curved segment is observed when *C*_i_ is ~200 μmol·mol^−1^ to *CSP*, and *P*_n_ increases slowly as *C*_i_ increases, gradually entering the restriction stage of RuBP regeneration[62]. Third, an almost linear segment when *C*_i_ > *CSP*, *P*_n_ changes insignificantly as *C*_i_ increases, moving into the restriction stage of triose-phosphate utilization (TPU). *P*_n_ at this stage is *P*_nmax_, which reflects photosynthetic electron transport and photophosphorylation activity [63].

The form of the *P*_n_-*C*_i_ curve changes when plants encounter stressful conditions, such as drought. Bernacchi *et al*.[64]considered that numerous factors would influence the curve of *P*_n_-*C*_i_ which included physiological changes (e.g. *V*cmax,*J*max or *R*d) and environmental changes (e.g. drought, temperature and/or atmospheric CO_2_ concentration). However, the quantitative relationship between this change and soil moisture has remained unclear. This study demonstrated that the photosynthetic *P*_n_-*C*_i_ curve of *H. rhamnoides* L. exhibited a classical form, with *P*_nmax_, *CE*, *CSP*, and *R*_p_ being high and *Γ* being low within a suitable *RWC* range (i.e., 42.8%−83.2%); *P*_n_ levels were highest when *RWC* was 71.3%(Fig. 1; Table 1). Three photosynthetic parameters, *P*_nmax_, *CE*, and *CSP*, declined dramatically when soil moisture was beyond this range. *H. rhamnoides* L. exhibited wide photosynthetic adaptability to soil moisture compared to the suitable *RWC* ranges of *Robinia pseudoacacia* L. (50.0%–81.6%), *Platycladus orientalis* L. (5.3%-75.0%)[65], *Syringa oblata* Lindl. (58.8%-76.6%) [66], and *Ziziphus jujube* (46.0%-80.5%)[67].

The common method for obtaining *CE* is the traditional linear regressive method, whereby *CE* is the slope of the straight line of the *P*_n_-*C*_i_ curve at a low CO_2_ concentration (*C*_i_ ≤ 200 μmol mol^− 1^)[35,68]. Many studies have shown that the *CE* values of different plants vary greatly[69,70]. For example, under normal growth conditions, the *CE* values of *Rheum tanguticum*, *Anisodus tanguticus*, and *Gentiana straminea* were approximately 0.0453, 0.1116, and 0.0902 [71], those of two pepper (*Capsicum annuum* L.) cultivars were approximately 0.145 and 0.159[74] (Hu *et al*. 2008), that of *Zantedeschia aethiopica* was approximately 0.074 [72](Yiotis&Manetas 2010) and that of *Sophora moorcroftiana* was about0.03[70]. Although Hu et al. [44]showed that soil moisture greatly affects the *CE* values of plants, the quantitative relationship between *CE* and soil moisture has remained unclear. According to a previous study, the *P*_n_-*C*_i_ curve of photosynthesis does not have a strictly linear relationship at a low CO_2_ concentration [43].

### CO_2_ response curves and photosynthetic parameters fitted by different models

The major use of different CO_2_ response models lies in the equations used to fit the CO_2_ response and its characteristic parameters to extract physiologically meaningful variables; these parameters are used to describe physiological responses of leaves to different treatments [64,73]. For example, *CE*_Γ_ at the CO_2_ compensation point, *CE*_0_, and the absolute value of the slope of the line between *C*_i_ = 0 and *C*_i_ = *Γ* on the *CE*_Γ0_ curve can be fitted, and they have clear physiological meaning and unique values. However, the applicability and simulated accuracy of the empirical models are limited by their asymptotic form with no extreme values[38,39] (Ye & Gao 2009, Ye 2010). In some studies[43,45,46], *P*_nmax_ was much larger than the measured value, while *CSP* was far less than the measured value. In particular, the CO_2_ response curves could not be fitted under stressful conditions. The same problem was noted in this study.

Although the modified rectangular hyperbola model proposed in recent years can fit and analyze various forms of CO_2_ response curves more accurately[77,41], overcoming the limitations of other models to a certain extent, there are few reports regarding its application in plants under different soil moisture conditions. This study indicated that when the soil moisture was within a suitable *RWC* range, the CO_2_ response curves and characteristic parameters were well fitted by the four models (*R*^2^ > 0.99,Fig. 2; Table 2), where the non-rectangular hyperbola model and modified rectangular hyperbola model fit the data better than the other two models(Fig. 2 B,D). When soil moisture was too high or too low, the modified rectangular hyperbola model was better than the other three models fitting the CO_2_ response process and its characteristic parameters in the leaves of *H.rhamnoides* L.(Fig. 2 D). This result is consistent with the findings of Jiao & Wei [45]and Lv *et al.*[46]. This study demonstrated that the simulation results of the photosynthetic-CO_2_ response model were closely related to soil moisture content.

## Conclusions

Research on the effects of soil moisture on the physiological mechanisms related to photosynthetic responses is garnering attention toward CO_2_ response curves and photosynthetic parameters in trees. This study indicated that soil moisture content affected the CO_2_ response processes in the leaves of *H.rhamnoides* L. The photosynthetic *P*_n_-*C*_i_ curve exhibited a classical form, with *P*_nmax_, *CE*, *CSP*, and *R*_p_ being high, while *Γ* was low when the *RWC* ranged from 42.8% to 83.2%. *H.rhamnoides* L. exhibited high photosynthetic efficiency in this soil moisture range, and the *P*_n_ levels were highest when *RWC* was 71.5%. Three photosynthetic parameters, *P*_nmax_, *CE*, and *CSP*, declined dramatically when soil moisture was outside the aforementioned range. Thus, the suitable *RWC* for *P. sibirica* L. ranged from 46.5% to 81.6%, and the most suitable *RWC* was ~66.8%.

The *CE* (i.e *CE*_0,_*CE*_Γ_,*CE*_Γ0_)values of *H. rhamnoides* L. were significantly different under different soil moisture conditions. For example,the *H. rhamnoides* L. *CE*_Γ0_ ranged from 0.0260 to 0.0564, with a comparatively higher level > 0.047 in the *RWC* range of 42.8%-83.2%; the maximum (0.0564) appeared when *RWC* was71.5%. *CE* of *H. rhamnoides* L. decreased markedly when the soil moisture was too high or too low. When soil moisture was within the suitable *RWC* range, the CO_2_ response curves and characteristic parameters were well fitted by the four models (*R*^2^ > 0.99). The non-rectangular hyperbola model and modified rectangular hyperbola model fitted better than the other two models (*R*^2^ > 0.998). However, when soil moisture exceeded the suitable *RWC* range, only the modified rectangular hyperbola model fit the CO_2_ response curves and photosynthetic parameters well. Compared to the other three models, the modified rectangular hyperbola model demonstrated extensive applicability for fitting photosynthetic CO_2_ responses under different soil moisture conditions.

## Acknowledgments

This work was funded by the Doctoral Research Fund of Shandong Jianzhu University (No. XNBS1420). We thank LetPub (www.letpub.com) for its linguistic assistance during the preparation of this manuscript.

## Name and abbreviation

*RWC*: relative soil water content
*MWC*: soil mass water content
*FC*: field water capacity
*P*_*n*_: net photosynthetic rate
*C*_*i*_: intercellular CO_2_ concentration
*CSP*: CO_2_ saturation point
*Г*: CO_2_ compensation points
*CE*: carboxylation efficiency
*R*_*p*_: photorespiration rate

